# A bioactive peptide from the pearl has dual roles in resisting SARS-CoV-2 infection and its complications

**DOI:** 10.1101/2023.10.23.563427

**Authors:** Xiaojun Liu, Yayu Wang, Zehui Yin, Qin Wang, Xinjiani Chen, Bailei Li, Liping Yao, Zhen Zhang, Rongqing Zhang

## Abstract

Angiotensin-converting enzyme 2 (ACE2) is a critical receptor for the entry of the SARS-CoV-2 virus into cells. Moreover, a decrease in ACE2 level and its activity due to SARS-CoV-2 infection is considered a crucial reason for the development of Covid-19-associated complications. Here, we report a bioactive peptide derived from the seawater pearl oyster *Pinctada fucata*, named SCOL polypeptide, which binds strongly to ACE2 and effectively inhibits 65% of the binding of the SARS-CoV-2 S protein to ACE2; thus, this peptide can be used as a blocker to enable cells to resist SARS-CoV-2 infection. The SCOL polypeptide also increases ACE2 enzyme activity by 3.76 times. Previous studies have shown that ACE2 deficiency is associated with inflammation, pain, cardiovascular diseases, insulin resistance, and nervous system injury. Therefore, the SCOL polypeptide can be used to treat or alleviate complications such as lung inflammation, pain, diabetes, cardiovascular diseases, and loss of taste or smell caused by SARS-CoV-2 infection. Thus, the SCOL polypeptide can play a dual role in resisting SARS-CoV-2 infection.

## Introduction

Severe acute respiratory syndrome coronavirus 2 (SARS-CoV-2), the causative agent of the global coronavirus disease 2019 (Covid-19) pandemic in 2020, is the third novel coronavirus to cause a life-threatening pneumonia pandemic in humans, along with SARS-CoV and Middle East respiratory syndrome virus (MERS-CoV). One of the key factors that enable these viruses to overcome the immune barrier is their spike (S) protein that can effectively bind to angiotensin-converting enzyme 2 (ACE2) on the human cell surface and mediate the entry of the virus into the cells [1, 2]. The S protein contains two functional subunits, namely S1 and S2. The first step of virus entry into the cells is the binding of the N-terminal of the S1 subunit to the pocket structure of ACE2; the S2 subunit then mediates the fusion of the virus with the cell membrane [1, 3]. Thus, ACE2 functions as a gateway for the entry of the SARS-CoV-2 virus into the cells, and the distribution of this receptor in the body determines the susceptible sites of SARS-CoV-2 infection.

ACE2 is mainly expressed in the heart, intestine, lung, kidney, testis, brain, and other tissues [4-7]. Because ACE2 is mostly present as a cell membrane receptor in these tissues, the SARS-CoV-2 virus can easily enter the human body through the epithelial cells of these tissues; in particular, the wide surface area of alveolar epithelial cells facilitates the easy entry of the virus, resulting in pneumonia as the primary symptom of SARS-CoV-2 infection [7].

ACE2 is a type I transmembrane glycoprotein with a single extracellular catalytic domain. ACE2 can convert angiotensin I into angiotensin 1-9 and balance the action of ACE by reducing the substrate level. Angiotensin II (Ang II) is converted into angiotensin 1-7 (Ang-(1-7)) by ACE2, and Ang-(1-7) plays a physiological role by binding to its receptor Mas [8]. Ang-(1-7) can induce vasodilation and decrease blood pressure, and it can be used as a beneficial vasodilator and anti-proliferative agent [9, 10]. Therefore, ACE2 plays a role as a stabilizer in the ACE1/AngII/AT1 receptor axis and the ACE2/Ang1-7/MAS receptor axis, the two major pathways in the renin–angiotensin system (RAS). Many complications caused by SARS-CoV-2 infection are related to ACE2 deficiency during virus invasion [7].

The loss of ACE2 can lead to the accumulation of Ang II in the body, resulting in a series of adverse reactions, including myocardial hypertrophy and dysfunction, interstitium fibrosis, endothelial dysfunction, increased inflammation and oxidative stress, and increased coagulation [7, 11-13]. Lung inflammation caused by SARS-CoV-2 infection and acute respiratory distress syndrome (ARDS) are partly associated with this phenomenon. It has also been confirmed that Ang II can interfere with the adaptive immunity of the body’s immune system and increase the number of inflammatory factors, which further reduces the body’s resistance to SARS-CoV-2 invasion [14, 15]. Therefore, it is crucial to develop viral blockers targeting the common gateway through which the coronavirus enters human cells. Moreover, these blockers could be considered worthwhile if they do not affect ACE2 activity or even enhance it.

In the present study, we describe a bioactive peptide (SCOL polypeptide) derived from the seawater pearls of *Pinctada fucata*. We found that this bioactive peptide has the potential to serve as a blocker for SARS-CoV-2 infection.

## Results

### 1. Isolation and identification of SCOL polypeptide

We first extracted the pearl matrix protein from the nucleus-free pearl of *Pinctada fucata* by decalcification. The extracted pearl matrix protein was then hydrolyzed using trypsin. The enzyme-digested pearl matrix protein-polypeptide mixture was separated by high-performance liquid chromatography, and samples with a retention time of 9.5 min were collected. The collected fraction had a molecular weight of 1794.8 Da, and its amino acid sequence was IPSTTPFPSTTVATTTM (Figure 1A), as determined by matrix-assisted laser desorption ionization time-of-flight mass spectrometry (MALDI-TOF/TOF-MS).

**Figure 1.**
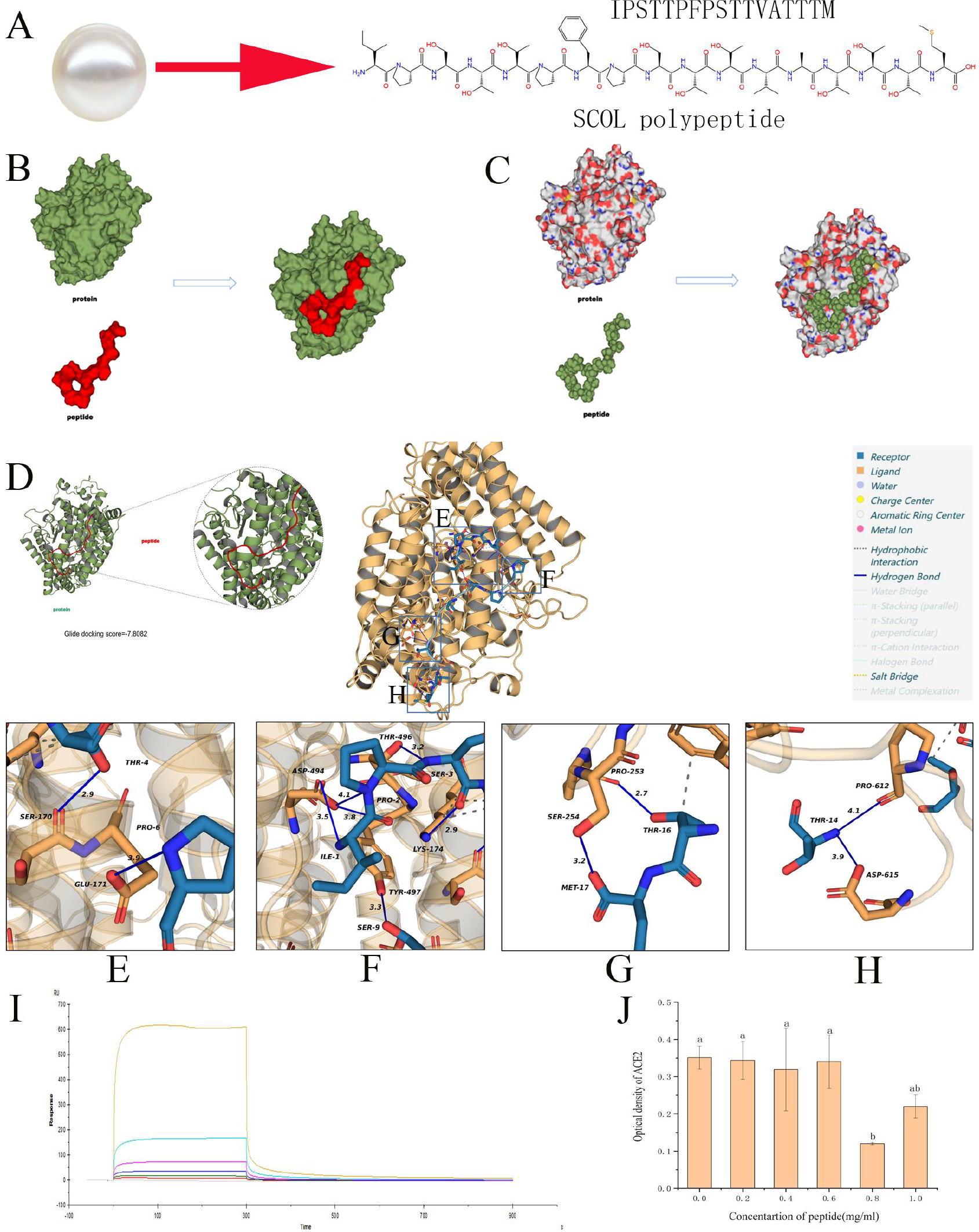
The SCOL polypeptide and its characters. A, the SCOL polypeptide with the amino acid sequence “IPSTTPFPSTTVATTTM” isolated from pearls of *P. fucata*; B-H, molecular docking results between the SCOL polypeptide and the ACE2 protein; I, binding ability of SCOL polypeptide to ACE2 detected by Biacore X100; J, Competitive binding analysis of SCOL polypeptide and the S protein to ACE2, the highest inhibitory rate of the S protein binding to ACE2 inhibited by SCOL polypeptide was 65.8%.

### 2. Molecular docking of SCOL polypeptide with ACE2

Figure 1 shows the interaction between the protein and the polypeptide analyzed by the Protein-Ligand Interaction Analyzer (PLIP) server by using the protein as a reference chain for the interaction analysis, with gold and blue representing the protein and polypeptide, respectively (Figure 1 B-H). As shown in Figure 1E, hydrogen bonds are formed between Pro-6 of the polypeptide and Glu-171 of the protein (solid blue line) and between Thr-4 of the polypeptide and Ser-170 of the protein. Figure 1F shows the formation of hydrogen bonds between Asp-494 of the protein and Ile-1 and Pro-2 residues of the polypeptide, and between Tyr-497 of the protein and Ser-9 of the polypeptide. As shown in Figure 1G, hydrogen bonds are formed between Met-17 of the polypeptide and Ser-254 of the protein, and between Pro-253 of the protein and Thr-16 of the polypeptide. Figure 1H shows the formation of hydrogen bonds between Asp-615 and Pro-612 residues of the protein and Thr-14 of the polypeptide. Several sets of hydrophobic interactions (gray dashed line) are also noted.

### 3. Binding ability of SCOL polypeptide to ACE2

Biacore X100 was used to measure the binding ability of SCOL polypeptide to ACE2. The ACE2 protein was set in a chip, and a concentration gradient of SCOL polypeptide was then added to mobile phase. The test results are shown in Figure 2. The test curves of SCOL polypeptide at 100, 50, 25, 12.5, 6.25, and 3.125 μg/mL concentrations are shown from top to bottom (Figure 1I), respectively. The dissociation constant KD (M) value was 7.687 × 10^-5^, thus indicating that SCOL polypeptide has a high affinity for ACE2.

**Figure 2.**
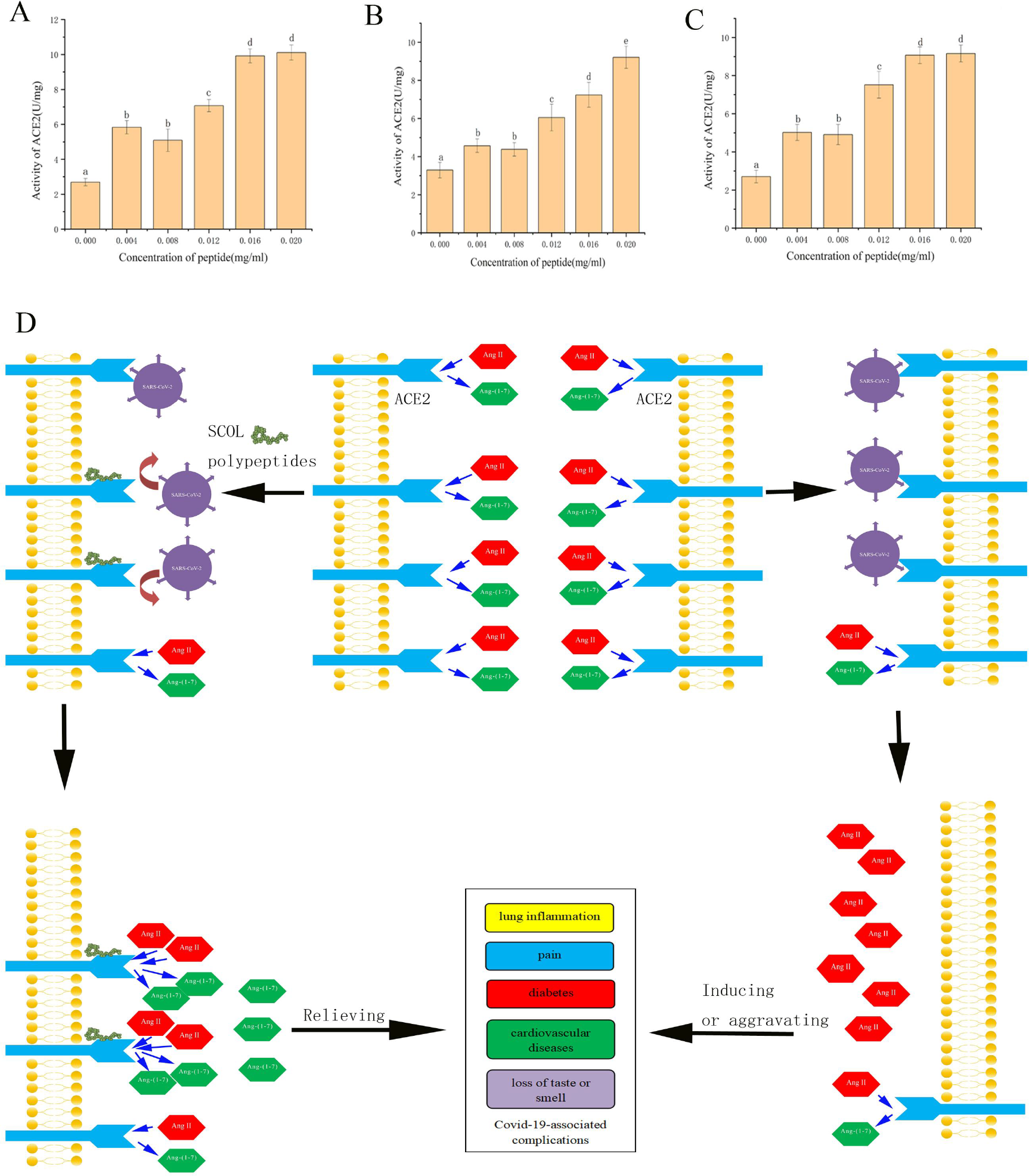
Increasing the enzyme activity of ACE2 by SCOL polypeptide and the significance in resisting SARS-CoV-2 infection. Effects of SCOL polypeptide on ACE2 enzyme activity in rat liver (A), kidney (B), and heart (C). D, the significance of SCOL polypeptide in resisting SARS-CoV-2 infection .

### 4. Competitive binding analysis of SCOL polypeptide and the S protein to ACE2

The RayBio®Covid-19 Spike-ACE2 binding assay kit was used to detect the binding of the S protein and SCOL polypeptide to ACE2. The recombinant S protein was bound to the 96-well plates in the kit, and the polypeptide was then added to the wells in the presence of ACE2. Small molecular peptides bound to the key sites of ACE2, thereby reducing the rate of binding of the S protein to ACE2. Compared to the control group, SCOL polypeptide at all concentrations showed inhibitory effects, and the highest inhibitory rate was 65.8% for the 0.8 mg/mL group (Figure 1J), thus indicating that SCOL polypeptide exhibits a strong binding ability to ACE2 and can competitively inhibit the binding of the novel coronavirus S protein to ACE2. Thus, SCOL polypeptide has a high potential as a competitive inhibitor of the novel coronavirus S protein.

### 5. Effects of SCOL polypeptide on ACE2 enzyme activity

The ACE2 active fluorescence assay kit (Biyuntian) was used to detect the effect of SCOL polypeptide on ACE2 enzyme activity. The assay is based on the following principle. In the absence of the ACE2 enzyme-substrate reaction, the substrate remains intact. Consequently, the two fluorophores are close enough and undergo fluorescence resonance energy transfer, resulting in no fluorescence detection. In contrast, following the enzyme-substrate reaction, the substrate is degraded by ACE2, and the ends of the polypeptide are separated. Consequently, the two fluorophores are separated, and fluorescence can be detected. Thus, the enzyme activity of ACE2 can be determined very sensitively through fluorescence detection. The assay showed that SCOL polypeptide increases the activity of the ACE2 enzyme in the liver, kidney, and heart of mice by up to 3.76 (Figure 2A), 2.8 (Figure 2B), and 3.38 (Figure 2C) times that in the tissues without the polypeptide, respectively.

## Discussion

In the present study, we observed that SCOL polypeptide can effectively bind to ACE2. The results of molecular docking showed the formation of hydrogen bonds between the Ile-1, Pro-2, Thr-4, Pro-6, Ser-9, Thr-14, Thr-16, and Met-17 residues of the polypeptide and Asp-494, Ser-170, and Glu-171 residues of the ACE2 protein. Tyr-497, Pro-253, Asp-615, and Pro-61 formed hydrogen bonds, and several hydrophobic interactions between the SARS-CoV-2 S protein and ACE2. Biacore X100 was used to measure the binding ability of SCOL polypeptide to ACE2. The results showed that SCOL polypeptide exhibits a strong affinity for ACE2. Because ACE2 functions as a receptor for SARS-CoV-2, we examined whether SCOL polypeptide and the S1 subunit of the SARS-CoV-2 S protein compete to bind to ACE2; this is a key aspect to understand whether SCOL polypeptide can act as a blocker for cell invasion by the SARS-CoV-2 virus. Further experiments showed that the peptides in SCOL polypeptide competitively prevented the binding of the SARS-CoV-2 S protein to ACE2 (up to 65.8%). The experiments revealed the potential of SCOL polypeptide to function as a blocker of cell invasion by the SARS-CoV-2 virus, and further studies are required to understand the specific mechanisms involved in the competitive binding to ACE2 between SCOL polypeptide and the S1 subunit of the SARS-CoV-2 S protein.

SCOL peptide can also increase the activity of the ACE2 enzyme. Following cell invasion by the SARS-CoV-2 virus, the virus binds to the ACE2 receptor on the cell membrane to form a complex. The fusion of the virus with the cell membrane allows its entry into the cell, leading to the downregulation of the receptor and resulting in the loss of ACE2. Consequently, the ACE2 enzyme activity on the cell membrane is reduced; however, it remains unclear whether the enzyme is completely inactivated [2, 16, 17]. Therefore, the invasion of the SARS-CoV-2 virus into the cells may decrease ACE2 enzyme activity and function. As mentioned previously, the downregulation of ACE2 activity leads to Ang II accumulation and Ang-(1-7) decrease in tissues or cells, which may contribute to induce or exacerbate some Covid-19-related symptoms. The underlying mechanisms associated with the main symptoms of Covid-19 are as follows.

AngII is considered a major factor in the release of inflammatory cytokine mediators [18]. According to previous studies, AngII may mediate the development of inflammatory lesions associated with SARS-CoV or acute lung injury in the respiratory system (alveolar wall thickening, edema, etc.) [19, 20]. Ang-(1-7) can inhibit inflammation by inhibiting the NF-kB pathway [21], thus playing an opposite role to AngII. Lung inflammation and the resulting ARDS are potentially fatal complications of SARS-CoV-2 infection, and the downregulation of ACE2 reduces the amount of Ang-(1-7) and accelerates AngII accumulation, thus enhancing the deteriorative effect of the disease. In clinical studies on the treatment of pneumonia, Ang-(1-7) was found to play an anti-inflammatory role by reducing perivascular and peribronchial inflammation and by preventing or relieving pulmonary fibrosis after inflammation [21-24].

Hypertension and related cardiovascular diseases are also common complications in Covid-19 patients [25, 26]. Blood pressure in humans is maintained at a normal level through the balance between the ACE1/Ang II/AT1 receptor axis and the ACE2/Ang-(1-7)/MAS receptor axis, which are the two major pathways of the RAS. SARS-CoV-2 infection can induce the downregulation of ACE2 and thus increase Ang II levels. Ang II accumulation in the body can lead to vasoconstriction and increased blood pressure, and this could further worsen cardiovascular diseases such as coronary heart disease, heart failure, arrhythmia, and thrombosis [27, 28]. Previous studies on rats have shown that Ang-(1-7) can significantly reduce various treatment-induced arrhythmias [29], cardiac fibrosis [30], myocardial hypertrophy [31], and cardiac hypertrophy [29]. It also prevents cardiac dysfunction and left ventricular remodeling [32]. Ang-(1-7) can also play an antithrombotic role through MAS receptors [33]. Therefore, the increase in ACE2 activity may contribute to the beneficial role of the ACE2/Ang-(1-7)/MAS receptor axis in cardiovascular diseases.

Diabetes is also a common complication in Covid-19 patients [25, 26]. Ang II interferes with the PI3K/AKT signaling pathway, an important pathway of insulin signal transduction; suppresses the protective effects of insulin on blood vessels, and enhances insulin resistance through AT1 receptors [34]. Ang-(1-7) can reduce insulin resistance by activating GSK3, a downstream target of the PI3K/AKT signaling pathway [35, 36]. In a rat model, the increase in the Ang-(1-7) level at the normal food intake level was found to enhance insulin sensitivity and insulin-stimulated glucose uptake [37]. Therefore, the increased expression or activity of ACE2 is critical for treating diabetes.

Headache, myalgia, and chest pain are the most common types of pain experienced by Covid-19 patients [38, 39]. Pain is a nonspecific symptom of Covid-19 patients. Although the mechanism of pain caused by SARS-CoV-2 infection is complex, ACE2 loss could be one of the causes. Ang II can activate p38 MAPK through the AT1 receptor to produce intense pain behavior in mice [40]. For example, it is suspected that the virus invasion of the spinal cord with neurons and microglia leads to a decrease in ACE2 levels, Ang II accumulation, and Ang-(1-7) levels in the spinal dorsal horn of Covid-19 patients, and this ACE/ACE2 imbalance will eventually induce pain in the spinal dorsal horn [41]. A previous study noted that the intrathecal administration of Ang-(1-7) in the spinal cord of mice inhibited Ang II-induced pain through MAS1 receptors [42].

As shown by previous studies, the SARS-CoV-2 virus invades human cells with neuronal properties, thereby resulting in a range of nonspecific disorders such as loss of smell and taste, cognitive and language disorders, and headaches, including those described above [43]. In addition to excessive inflammation, oxidative stress can also cause neurological dysfunction [43]. Elevated Ang II levels are considered the main cause of the release of reactive oxygen species and inflammatory mediators, which induce oxidative stress through the activation of NADH oxidase; this oxidative stress can damage cells and lead to neuronal degeneration [18]. As mentioned above, ACE2 can play an anti-inflammatory role through Ang-(1-7). However, the mechanism through which ACE2 participates in the antioxidant process is unclear. The loss of smell and taste is also related to the balance of hydrolyzed electrolytes in smell and taste cells, wherein the RAS system plays a key role. ACE2 downregulation caused by SARS-CoV-2 infection will disrupt this balance; the specific mechanism responsible for this effect requires further research [44, 45].

In light of these findings, scientists often face a dilemma when developing drugs to prevent and treat Covid-19. It might be feasible to restrict the entry of the SARS-CoV-2 virus into cells by reducing ACE2 expression with drugs; however, this approach often increases the imbalance of the RAS system, which may aggravate the complications caused by the SARS-CoV-2 virus. Some studies have shown that ACE inhibitors and Ang II receptor blockers (ARBs) used for treating Covid-19 complications can induce increased ACE2 expression [46-48] and accelerate virus spread.

New SARS-CoV-2 mutants continue to emerge globally, and vaccines developed against the original SARS-CoV-2 strain may be at risk of becoming less effective in the future. Moreover, epidemics caused by coronaviruses (such as SARS-CoV and MERS-CoV) have occurred several times in the past two decades, thus indicating the major threat posed by coronaviruses to humans. At present, it is impossible to predict whether new coronaviruses will cause pandemics in the future. The present study revealed that SCOL polypeptide can bind to ACE2. Although SCOL polypeptide can inhibit the binding of the SARS-CoV-2 S protein to ACE2 by only 65%, it can still prevent some loss of ACE2 due to virus invasion. More importantly, SCOL polypeptide increases the activity of ACE2 by more than 3-fold. ACE2 can efficiently catalyze Ang II, with an efficiency of approximately 400-fold that of Ang I [49]; thus, it can compensate for the loss of ACE2 in quantity to a certain extent (Figure 2D). Therefore, SCOL polypeptide has the potential to function as an effective anti-coronavirus drug.

## Materials and methods

### 1. Preparation of SCOL polypeptide

#### 1.1 Extraction of the pearl matrix protein

Seedless pearls of *P. fucata* from Zhanjiang, Guangdong Province, were washed, dried, and completely crushed with a grinder. The pearl powder was collected and placed in a beaker; next, 1.5 M EDTA was added at five times the volume of the powder, and the powder was stirred and decalcified in a 4°C chromatographic freezer. After 24 h of decalcification, the powder-EDTA mixture was placed in a 50 mL high-speed centrifuge tube for centrifugation at 10,000 rpm. The supernatant was collected, pumped, filtered, dialyzed, and freeze-dried to obtain the pearl matrix protein.

#### 1.2. Enzymatic hydrolysis of the pearl matrix protein

The lyophilized pearl matrix protein powder was obtained, and trypsin was added to the powder for enzymolysis. The activity of trypsin was 10,000 U/g, and the amount of trypsin added was 5% of the weight of the pearl matrix protein. The pH of the enzymolysis system was adjusted to 8.0, and enzymolysis was conducted in a water bath at 50°C for 2 h. The enzymolysis process was then inactivated by placing the mixture in a boiling water bath. After centrifugation at 10,000 rpm, the supernatant was collected. Following extraction and filtration, Millipore centrifuge tubes were used for centrifugal ultrafiltration (4000 rpm). The embedded Ultracel-PL ultrafiltration membrane with low adsorption was used to retain the fraction with a molecular weight of 3 kDa. The filtered solution containing the fraction with a molecular weight of less than 3 kDa was collected, concentrated, and freeze-dried to obtain the enzymatic pearl matrix protein-polypeptide mixture.

#### 1.3 Isolation and purification of the pearl matrix protein-polypeptide mixture

The enzymolized pearl matrix protein-polypeptide mixture was purified by high-performance liquid chromatography using a C18 column with mobile phase A consisting of deionized water containing 0.1% trifluoroacetic acid and mobile phase B consisting of acetonitrile containing 0.1% trifluoroacetic acid. The UV detection wavelength was 280 nm, the flow rate was 1 mL/min, and samples with a retention time of 9.5 min were collected. The target polypeptide was obtained by freeze-drying after concentration.

#### 1.4 Sequence analysis of SCOL polypeptide

SCOL polypeptide derived from the pearl matrix protein was determined by MALDI-TOF/TOF-MS. The molecular weight of the polypeptide was 1794.8 Da, and its amino acid sequence was as follows:

Ile-Pro-Ser-Thr-Thr-Pro-Phe-Pro-Ser-Thr-Thr-Val-Ala-Thr-Thr-Thr-Met

### 2. Molecular docking of SCOL polypeptide with ACE2

The protein structure of ACE2 was downloaded from PubChem. The peptides were mapped and ionized using pyMOL 2.5. The molecular docking sites of SCOL polypeptide and ACE2 were determined using the PLIP server (https://plip-tool.biotec.tu-dresden.de/plip-web/plip/index).

### 3. Determination of the binding ability of SCOL polypeptide to ACE2

Biacore X100 was used to determine the binding capacity of SCOL polypeptide to ACE2. The recombinant human ACE2 protein was purchased from Abcam. The ACE2 protein was fixed on the surface of an S-series SA chip (GE, lot number: 20139043) with 10× HBS-EP+ buffer by the biotin-streptavidin method. SCOL polypeptide solutions with different concentrations, namely 3.125, 6.25, 12.5, 25, 50, and 100 μg/mL, were prepared. The experimental conditions were as follows: flow rate: 50 μL/min; binding time: 300 s; reaction temperature: 25°C; dissociation flow rate: 30 μL/min; and dissociation time: 600 s.

### 4. Competitive binding analysis of SCOL polypeptide and the S protein to ACE2

The RayBio®COVID-19 Spike-ACE2 binding assay kit was used for the experiment.

#### 4.1 SCOL polypeptide and ACE2 protein mixture preparation

The 100× ACE2 protein concentrate in the kit was diluted to a 10× ACE2 protein solution with a diluent, and the SCOL polypeptide was diluted to concentrations of 0.2, 0.4, 0.6, 0.8, and 1 mg/mL with the diluent. In a PCR tube, 110 μL of each concentration of SCOL polypeptide was added, followed by the addition of 11 μL of the 10× ACE2 protein solution with vortex mixing. A control group without SCOL polypeptide was prepared by mixing 110 μL of the diluent and 11 μL of the 10× ACE2 protein solution, and three replicates were used for each group.

#### 4.2 Detection of the binding rate of the ACE2 protein and S protein by ELISA

The bottom of a 96-well plate was coated with the S protein binding domain. Next, 100 μL solution was drawn from the PCR tube of each group described in step 4.1 and added to the 96-well plate by using a row gun. The plate was sealed with a plate sealing film and incubated overnight at 4°C with gentle shaking. The supernatant was discarded, and the plate was washed 4 times with a washing solution; 300 μL of the washing solution was added to each well for cleaning. Next, 100 μL of 1× goat-derived anti-ACE2 antibodies capable of binding to the S protein-ACE2 protein complex was added to each well; the plate was gently shaken, incubated at room temperature for 1 h, and then washed with the washing solution. Subsequently, 100 μL of 1× HRP-labeled anti-goat-LGG antibodies was added to each well; the plate was gently shaken and incubated at room temperature for 1 h, followed by washing with the washing solution. Next, 100 μL of the color development agent TMB was added to each well, and the plate was incubated at room temperature in the dark for 30 min. The reaction was terminated by adding a termination solution, and the absorbance was measured at 450 nm. The absorbance reading was proportional to the amount of the S protein-ACE2 protein complex.

### 5. Determination of the effect of SCOL polypeptide on ACE2 activity

The ACE2 active fluorescence assay kit was used for the experiment. Briefly, 50 mg mouse tissue (liver, kidney, and heart) was collected and homogenized with 500 μL lysate; the mixture was then centrifuged at 12,000 × *g* at 4ºC for 5 min, and the supernatant was used for the next step. To obtain a standard curve, the standard solution was diluted with a buffer to yield a concentration of 0, 0.25, 0.5, 1, 2, 3, 4, and 5 μM. Next, 100 μL of the diluted standard solution was added to each well in triplicate to obtain the standard curve. Subsequently, 2 μL substrate was added to the 96-well plate. 2.2 μL SCOL polypeptide was added to the PCR tube at concentrations of 0.004, 0.008, 0.012, 0.016, and 0.02 μg/μL in the total system. Then the superserum sample of 8.8 μL in step (1) was added, and the buffer solution of 96.8 μL was added. Next, 11 μL of lysate and 96.8 μL of buffer were added into the blank control, mixed evenly, and 98 μL of the above solution was absorbed by a volute and added into the 96-well plate containing the substrate. Following the mixing of the solutions, fluorescence detection was performed with a fluorescent spectrometer. The detection was conducted at 37°C, with excitation and emission wavelengths of 325 and 393 nm, respectively.

The ACE2 protease activity was determined as follows:

ACE2 activity = A × V × T × C (U/mg),

where A represents the product in the sample; V is the sample volume; T is the time interval for the fluorescence intensity of the sample to show a linear relationship, and the change in the fluorescence intensity in T is measured as ΔRFU; and C is the sample protein concentration.

## Acknowledgments

This work was financially supported by the National Natural Science Foundation of China (32072975).

